# True Digestibility and Growth Response of Coated Essential Amino Acid Supplementation in Plant-Based Feeds for Rainbow Trout (*Oncorhynchus mykiss*)

**DOI:** 10.1101/2025.10.21.683664

**Authors:** Simon. J. Davies, Sadia Nazir, Matt E. Bell

**Affiliations:** School of Natural Sciences, University of Galway, Ireland; Department of Fisheries and Aquaculture, University of Veterinary and Animal Sciences, Lahore, Pakistan; School of Biological and Marine Sciences, University of Plymouth, UK

**Keywords:** Rainbow trout, plant proteins, crystalline amino acids, coated supplementation, true digestibility of EAAs, growth, feed utilisation

## Abstract

In order to assess the efficacy of using essential amino acids to support the inclusion of high plant based proteins in diets for rainbow trout, a 56-day growth and feed utilisation study was conducted with juvenile (60g) rainbow trout (*Oncorhynchus mykiss*) under controlled conditions. Fish were fed diets where a predominately white fishmeal based control diet was compared to diets where full fat soyabean meal (FFS) as well as maize gluten meal (MGM) were incorporated respectively at significant levels (up to 75% of the fishmeal). Ten essential amino acids (EAAs) supplemented these diets in both standard and coated forms to evaluate their potential to rebalance the dietary EAA profiles more precisely. All diets were adjusted to be isonitrogenous and isolipidic. In addition to the growth trial, a low purified protein diet was fed to the fish separately (at a mean weight of 200g) to collect faeces for determining endogenous protein loss as well as the individual experimental diets described. As such it was possible to calculate the true digestibility for essential amino acids.

The results indicated that corrective amino acid fortification of high plant diets, especially MGM could significantly (P<0.05) improve trout performance and raise their nutritional value. This was confirmed by final body weight, specific growth rate, feed conversion ratio and protein utilisation values. The true digestibility of the EAAs were similar and the values of each digestible EAA but coated EAAs were slightly superior in their digestibility coefficients. Coated EAAs improved digestibility coefficients by 2–5% and FCR by 8% compared to crystalline forms and also improved their utilisation leading to better trout performance with respect to growth, feed conversion efficiency and Nitrogen (crude protein) retention (P<0.05). The findings are discussed in the context of sustainable feeds and the circular bioeconomy for fish farming.

## Introduction

The aquaculture industry is increasingly focusing on sustainable and cost-effective feed formulations, particularly for species like rainbow trout (*Oncorhynchus mykiss*) and salmon (*Salmo salar*). As such, the global aquaculture industry faces increasing pressure to reduce its reliance on fishmeal (FM) as a primary protein source in aquafeeds due to economic and environmental concerns as reported by Sibley and Bell (2024). Traditionally, fishmeal has been the primary protein source in aquafeeds due to its high nutritional value. However, concerns over the environmental impact, sustainability, and rising costs of fishmeal have prompted research into alternative protein sources. Soybean meal (SBM) has emerged as a promising alternative due to its availability, cost-effectiveness, and favourable amino acid profile. However, SBM is deficient in certain essential amino acids (EAAs), such as methionine and lysine, and contains anti-nutritional factors that can impair growth performance and nutrient utilisation in carnivorous fish species like rainbow trout (Davies and Morris, 1997; Francis, Makkar and Becker, 2001). To address these limitations, supplementation with EAA mixtures has gained significant interest in enhancing growth rates and feed efficiency when SBM and other proteins are used as a partial or complete substitute for FM.

Plant-based ingredients, such as soybean meal and maize gluten, have emerged as promising substitutes. Soybean meal, in particular, has been extensively studied for its potential to replace fishmeal in rainbow trout diets. Research indicates that up to 50% of fishmeal can be replaced with enzyme-treated soybean meal without adverse effects on growth performance or feed conversion ratios in rainbow trout fry with considerations to Anti-Nutritional Factors (ANFs) in soybean meals as stated by Francis, Makkar and Becker (2001). Additionally maize gluten meal and various other legume, pulse and cereal derived protein concentrates have been utilised increasingly in compound aquaculture diets. However, replacing fishmeal with plant-based proteins can lead to imbalances in essential amino acids, which are crucial for fish growth and health. It has been stated previously that fish require a critical balance of essential amino acids (EAAs) rather than dietary protein *per se*. The combined EAAs constitute the ‘ideal protein’. This concept was first proposed by Wang and Fuller for pigs and is just as relevant for fish of all species (Wang and Fuller, 1989; van Milgen, and Dourmad, 2015). The ideal protein is where all 10 essential amino acids are rate limiting on their threshold requirements for growth and relative to lysine. In fish, this can often be related to the composition profile of muscle or gonadal tissue and the relative essential amino acid index.

To address this, supplementation of fish feeds with crystalline amino acids (CAAs) has been explored for many animal species and also with fish as described by Yamamoto, Sugita, and Furuita (2005). Studies have demonstrated that incorporating CAAs, such as lysine, methionine, and threonine, into plant-based diets can effectively improve growth performance and protein retention in rainbow trout to achieve optimal performance and better feed efficiency in production (Gaylord and Barrows, 2009).

Additionally, balancing amino acid profiles through CAA supplementation has been associated with improved weight gain and specific growth rates with better feed utilisation and nutrient retention. Replacing fishmeal with plant-based ingredients like soybean meal and maize gluten, supplemented with crystalline amino acids, offers a viable strategy for developing cost-effective and sustainable diets for many fish and more recently for rainbow trout (Gaylord and Barrows, 2009; Davies, Morris and Baker, 1997) and recently with tilapia (Davies and Bell, 2024). This approach not only maintains growth performance and feed utilisation but also contributes to the sustainability of aquaculture practices. Supplementation strategies typically involve the use of either crystalline or coated amino acids (in a protected shell of protein). Crystalline amino acids (CAAs) are free-form compounds that are rapidly absorbed in the gastrointestinal tract, often leading to mismatches in amino acid availability relative to protein synthesis needs. This asynchronous availability can reduce the efficiency of nitrogen utilisation and potentially limit growth performance (Davies and Bell, 2024; Green and Hardy, 2002). Conversely, coated amino acids (CoAAs) are encapsulated in protective layers of polymeric proteins designed to modulate their release and absorption rates, thereby aligning more closely with the temporal demands of protein synthesis as described by Zhou and Yue (2010). These differences in absorption dynamics suggest that CoAAs may outperform standard CAAs in sustaining optimal growth and feed conversion rates in fish.

Studies have shown that the effectiveness of EAA supplementation is influenced by factors such as dietary composition, fish size, and physiological state, as well as the specific amino acid delivery mechanism (Xing *et al*., 2024). Furthermore, understanding the bioavailability and metabolic fate of supplemented EAAs is crucial to optimizing their use in aquaculture feeds. In fact, delaying EAA absorption can act to enhance their efficiency to better time their uptake along with the complete protein degradation during the main digestion phase in the mid gut previously explained by Dabrowski and Dabrowska (1981). Salmonid species like trout and Atlantic salmon have quite stringent demands for balanced essential dietary amino acids and this becomes limiting when fishmeal is reduced at the expense of plant protein (Halver and Hardy, 2003). White fishmeal source is often used in many organic and sustainable fish production systems and currently an increasing use of lower specification fishmeal sources generated from trimmings and discards within the capture fisheries industry.

Consequently, the principal aims of this investigation were to evaluate the growth performance, feed utilisation, of rainbow trout fed soybean meal SBM and maize gluten meal based diets supplemented with either crystalline or coated EAA mixtures. By comparing these supplementation strategies, we provide insights into optimizing plant-based diets for sustainable aquaculture practices in lower white fishmeal type diet formulations. We aimed for integrative experimental objectives to evaluate the optimal use of essential amino acids in diets for rainbow trout incorporating high plant protein concentrates in a classical growth trial. A consecutive digestibility evaluation for the essential amino acids was also performed and an additional low protein diet formulated in order to ascertain their ‘true digestibility’ by quantitating the endogenous losses to correct apparent digestibility coefficient values.

## Material & Methods

### Diets

Six experimental diets were formulated for rainbow trout to test the feasibility of replacing the fishmeal component with either full fat soybean meal (FFS) or maize gluten meal (MGM) as major plant protein concentrates. The effect of using supplementary crystalline essential amino acids (CAAs and coated CoAAs) to restore dietary levels to those in the control white fishmeal and animal protein based diet were compared with EAA mixtures as well as coated (polymeric protein complex) versions for the plant substituted diets. All diets were formulated using FeedSoft™ (Austin, Texas) feed formulation software with a designated rainbow trout database for nutrient requirements. These experimental diets are presented in Table 1 showing their respective ingredient composition profiles.

**Table 1.** Composition of experimental diets fed to juvenile rainbow trout (*Oncorhynchus mykiss*) (g/kg)

| Ingredients | Ref diet | FFS | FFS+ EAAS | MGM | MGM+EAAS | MGM+EAAS coated |
| --- | --- | --- | --- | --- | --- | --- |
| White fishmeal (66) | 537.3 | 150 | 150 | 150 | 150 | 150 |
| Full-fat soya bean (36) | - | 566.3 | 555.5 | - | - | - |
| Maize gluten meal (60) | - | - | - | 283.3 | 235 | 235 |
| Poultry byproduct meal | 100 | 100 | 100 | 100 | 100 | 100 |
| Blood meal | 20 | 20 | 20 | 20 | 20 | 20 |
| Fish oil (Marine) | 81.8 | 15 | 15 |  |  |  |
| Wheat middling flour | 205.9 | 103.4 | 88.5 | 311.8 | 308.2 | 308.2 |
| Vitamin premix<sup>1</sup> | 20 | 20 | 20 | 20 | 20 | 20 |
| Mineral premix<sup>2</sup> | 20 | 20 | 20 | 20 | 20 | 20 |
| *Supplementary amino acids |  |  |  |  |  |  |
| D.L. Methionine | - | - | 5.4 | - | 4.8 | 4.8** |
| L Lysine | - | - | 5.8 | - | 15.2 | 15.2* |
| L Threonine | - | - | 3.0 | - | 5.5 | 5.5* |
| L Histidine | - | - | 0.7 | - | 2.4 | 2.4 |
| L Argine | - | - | 3.9 | - | 13.1 | 13.1 |
| L Isoleucine | - | - | 1.3 | - | 4.2 | 4.2 |
| L Leucine | - | - | 3.1 | - | - | - |
| L Valine | - | - | 2.9 | - | 4.4 | 4.4 |
| L Tryptophan | - | - | - | - | 1.3 | 1.3 |
| Dietary marker; chromic oxide (Cr<sub>2</sub>O<sub>3</sub>) | 5.0 | 5.0 | 5.0 | 5.0 | 5.0 | 5.0 |

1, 2, Vitamin and mineral premixes by PNP Ltd (Premier Nutrition Products) and supplying the NRC 2011 requirements for rainbow trout. \* Supplementary non-coated and coated amino acids were provided by Eurolysine, Avril Group, Amiens, France, AjiPro®-L; c-Lysine. \*\*DL-Methionine 50%, coating materials 50% Shandong Fulikang Animal Nutrition Co., Ltd. Shandong, China.

Diets were physically processed by blending the dry ingredients into a homogenous mixture with a Hobart A 120 industrial food extruder. The required supplementary oil for each diet was added gradually and after a few minutes of mixing, 350ml of water was added to prime the binder. Once a homogenous dough mixture was obtained, the diets were extruded through a mincer into ‘spaghetti-like’ strands and broken into smaller pellets. Pellets were dried by convection air in an oven at 70°C. After cooling, the diets were packed in sealed airtight containers and stored at 15°C until required. Prior to feeding the fish, the diets were further broken into smaller pellets (1-3mm diameter) to suit the gape size of juvenile and later production size rainbow trout for digestibility work.

### Fish and feeding trial regime

All female rainbow trout *(O. mykiss*) of mean weight 30g were randomly distributed (40 fish per tank) into each of six tanks in triplicate groups of 0.5m^3^ capacity within a semi-closed recirculation system with incorporated mechanical and biological filter compartments. The water temperature was held at a constant 15°C throughout the growth trial period of 8-weeks (56 days). A photoperiod of 12hrs light and 12hrs dark was maintained for the 8-week period. Water quality parameters were kept constant within limits tolerable to rainbow trout for ammonia, nitrite and nitrate with a 5% exchange per day with charcoal filtered mains water.

A feeding rate of 2% body weight per day was adopted and adjusted every two weeks on the basis of sample weight inventories. All rainbow trout were individually weighed at the start and end of the trial and bulk weighed at intermediate stages. On termination of the study, samples of fish (six per triplicate tanks) were taken for full proximate analysis to determine body composition. Rainbow trout were anaesthetised in buffered MS222 (200 mg/L), until loss of equilibria and response to human contact was observed. Subsequently, fish were subjected to lethal cranial percussion to ensure a humane endpoint.

### Digestibility trial and Faecal collection

In a separate trial for faecal collection, a selection of rainbow trout originating from the primary growth trial were allowed to attain a mean weight of 200g on a standard commercial trout diet (Biomar Ltd). Subsequently they were then returned to the original experimental diets for a four week acclimation period fed to apparent satiation, reaching a mean weight of 250g. During a one-week faecal sampling period, rainbow trout were anaesthetised in buffered MS222 (200 mg/L), until loss of equilibria and response to human contact was observed. Manual stripping of faeces was performed by hand, by lightly applying pressure to the hind portion of the abdomen. All twenty five fish per triplicate tank (25 fish per treatment group) were sampled as described. After rainbow trout were sampled, they were reintroduced to their respective tanks for full recovery and again sampled after a period of three days and further feeding. Faecal material was collected in aluminium trays over ice and pooled by tank number. The faecal samples collected were freeze-dried and homogenised with a pestle and mortar to achieve a fine texture. It should be noted that rainbow trout were fed the separate low protein diet for endogenous protein loss determination in the same manner described above for faecal sampling.

### Biometrics and calculations

From the mean initial and final weights and the values obtained for initial and final carcass protein content, Specific Growth Rate (SGR) and Apparent Net Protein Utilisation (NPU) for each dietary treatment could be calculated. Specific growth rate (SGR), the rate of growth of an animal, is a sensitive index of protein quality under controlled conditions being proportional to the supply of essential amino acids. It is the main growth assessment indicator for fish performance used in the scientific literature. Note: Mean Growth Rate (MWG); Average Daily Growth (ADG); Specific Growth Rate (SGR); Food Conversion Ratio (FCR); Apparent Net Protein Utilisation (ANPU); Protein Efficiency Ratio (PER). These were calculated according to the formulae presented below; Apparent Digestibility Coefficients (ADC) were determined using the formula listed below.

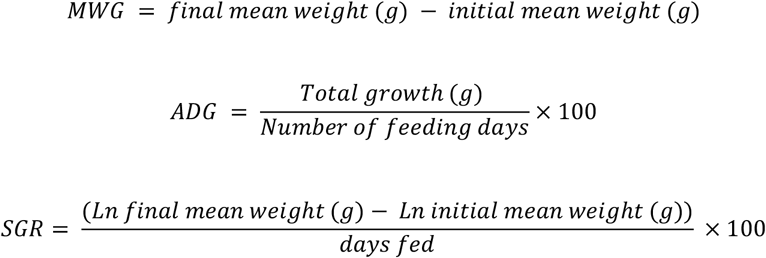

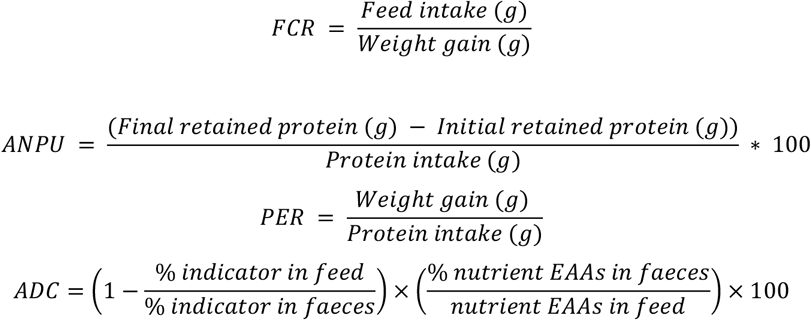

True protein and EAA digestibility were calculated using the formula above but based on corrections for the endogenous faecal N losses from feeding a low protein 2.00% diet as described in Table 2.

**Table 2.**
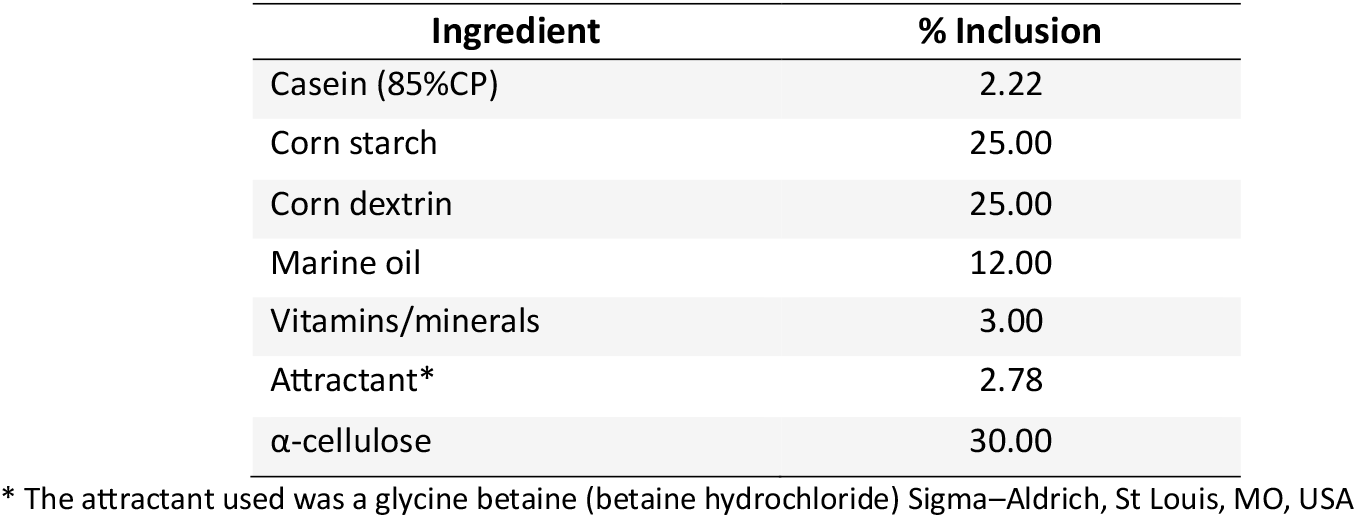
Formulation of a low protein 2.00% diet for rainbow trout (*Oncorhynchus mykiss*) for endogenous faecal nitrogen correction and amino acid digestibility studies.

| **Ingredient** | **% Inclusion** |
| --- | --- |
| Casein (85%CP) | 2.22 |
| Corn starch | 25.00 |
| Corn dextrin | 25.00 |
| Marine oil | 12.00 |
| Vitamins/minerals | 3.00 |
| Attractant* | 2.78 |
| α-cellulose | 30.00 |

\* The attractant used was a glycine betaine (betaine hydrochloride) Sigma–Aldrich, St Louis, MO, USA

### Ethics statement

The fish experimental procedures conformed to the European Community Directive (No. 2010/63/EU) (<u>EC, 2010</u>) and the UK Animal Scientific Procedures Act of 1986 (Home Office Licence PPL 30/2135). The investigations were approved by the local Institutional Animal Care Committee.

### Analytical methods and procedures

#### Determination of chromic oxide

Chromic oxide is the substance most commonly considered in evaluating the digestibility of experimental fish diets. The following method is a modification of the one proposed by Furukawa and Tsukahara (1966) so as to handle microsamples in determining the chromic oxide content of feed and faeces.

#### Proximate composition

Proximate analyses of diets (Table 3) and carcasses (Table 7) were made following the usual procedures (AOAC, 2019). These describe the classical Kjeldahl method for Crude Protein (N*6.25) and Soxtherm extraction method for Crude fat/oil extraction. Moisture and ash content of diets and fish were also determined by classical AOAC methods with slight modifications. These protocols were described in further detail for similar work conducted with rainbow trout (Green and Hardy, 2002).

**Table 3.** Nutrient analysis of experimental diets (%)

| **Nutrient** | **Ref diet** | **FFS** | **FFS+ EAAS** | **MGM** | **MGM+EAAS** | **MGM+EAAS coated** |
| --- | --- | --- | --- | --- | --- | --- |
| Moisture | 5.50 | 4.13 | 4.85 | 4.53 | 7.21 | 7.23 |
| Protein (Nx6.25) | 44.67 | 44.11 | 44.39 | 44.84 | 44.11 | 44.84 |
| Lipid | 12.00 | 13.73 | 13.50 | 12.00 | 12.50 | 11.92 |
| Ash | 16.67 | 12.45 | 12.67 | 11.06 | 10.56 | 9.95 |
| N.F.E* | 12.76 | 20.24 | 19.04 | 26.13 | 24.75 | 23.95 |
| Fibre | 2.11 | 3.83 | 3.67 | 2.68 | 2.62 | 2.58 |
| Calcium | 4.97 | 2.31 | 2.31 | 2.21 | 2.21 | 2.21 |
| Phos. | 2.63 | 1.48 | 1.46 | 1.40 | 1.39 | 1.39 |

\*Nitrogen Free Extractives (NFE) 100- [moisture+ crude protein+ ash+ lipid]
Nutrient analysis following standard AOAC methods for moisture, protein (N x 6.25; Kjeldahl), lipid, fibre and ash on ingredients and diet/fish tissue samples.

**Table 4.**
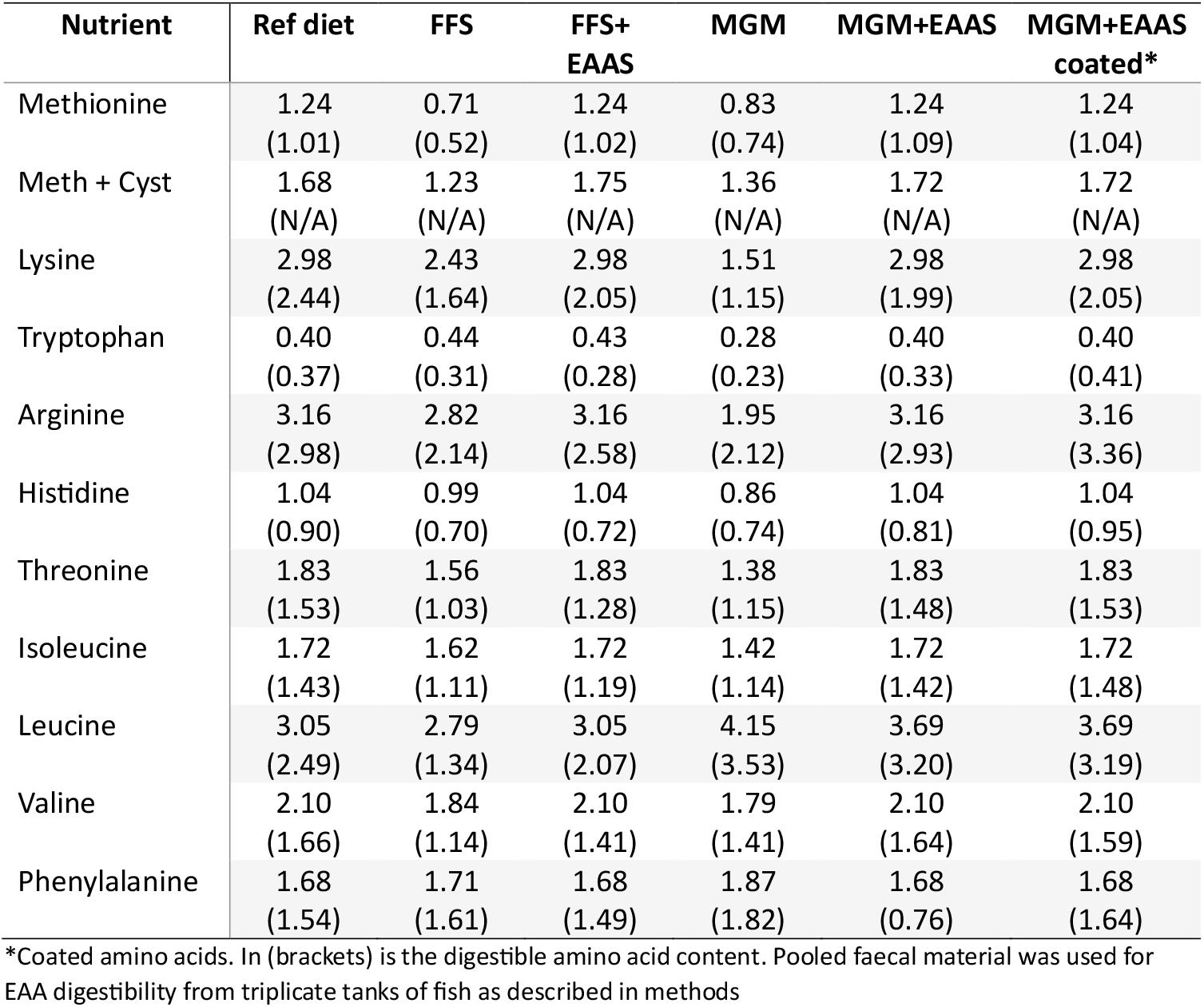
Essential and digestible amino acid profile of experimental diets (percent as fed)

**Table 5.** The coefficient of digestibility for individual essential amino acids for diets.

| Nutrient | Ref diet | FFS | FFS+ EAAS | MGM | MGM+EAAS | MGM+EAAS (coated) | SEM± |
| --- | --- | --- | --- | --- | --- | --- | --- |
| Methionine | 90.02<sup>a</sup> | 75.90<sup>c</sup> | 83.96<sup>b</sup> | 85.91<sup>b</sup> | 91.25<sup>a</sup> | 91.31<sup>a</sup> | 2.43 |
| Lysine | 89.14<sup>a</sup> | 75.17 | 75.47 | 77.61 | 85.83 | 87.13 | 2.58 |
| Tryptophan | 82.27<sup>b</sup> | 70.26<sup>c</sup> | 63.81<sup>d</sup> | 71.71<sup>c</sup> | 82.00<sup>b</sup> | 100<sup>a</sup> | 5.22 |
| Arginine | 88.80<sup>a</sup> | 79.97<sup>c</sup> | 81.13<sup>c</sup> | 85.25<sup>b</sup> | 89.61<sup>b</sup> | 92.85<sup>a</sup> | 2.07 |
| Histidine | 84.85<sup>a</sup> | 69.65<sup>d</sup> | 66.30<sup>d</sup> | 76.40<sup>c</sup> | 81.73<sup>b</sup> | 85.15<sup>a</sup> | 3.26 |
| Threonine | 88.46<sup>a</sup> | 71.35<sup>c</sup> | 72.43<sup>c</sup> | 82.36<sup>b</sup> | 86.65<sup>a</sup> | 88.56<sup>a</sup> | 3.22 |
| Isoleucine | 90.45<sup>a</sup> | 76.57<sup>c</sup> | 73.65<sup>c</sup> | 82.66<sup>b</sup> | 90.59<sup>a</sup> | 91.20<sup>a</sup> | 3.17 |
| Leucine | 88.55 | 52.05<sup>d</sup> | 70.55<sup>c</sup> | 87.24<sup>b</sup> | 90.43<sup>b</sup> | 90.65<sup>a</sup> | 6.37 |
| Valine | 87.87<sup>a</sup> | 69.83<sup>c</sup> | 70.83<sup>c</sup> | 82.12<sup>b</sup> | 86.98<sup>a</sup> | 81.58<sup>b</sup> | 3.19 |
| Phenylalanine | 100 | 100 | 100 | 100 | 100 | 100 | 0.00 |

Values are apparent true digestibility of individual amino acids +/- pooled standard error of mean. Values sharing the same superscripts are not significantly different (p<0.05)

#### Analysis of Amino acids in diets and faeces

Amino acids were analysed in feed ingredients and diets according to standard protocols for sample digestion and amino acid analyser protocols. The AA analyses of feed (ISO 13903-2005) were performed with a HPLC (Alliance System, Waters, Guyancourt, France) after hydrolysis of 100mg dried samples with 6 N HCl at 110°C for 23 h under reflux in small, sealed ampoules. Derivatisation with ninhydrin was done and by reaction with ninhydrin using spectrophotometric detection at 570 nm (440 nm for proline). This method has been the most widely performed method for several decades and is still commonly used. This method is official by Commission Regulation (EU) No 152/2009. IEC requires derivatization of AAs, which can be performed by post-column derivatization using ninhydrin. Post column ninhydrin derivatization has been the preferred method for many years by a majority of laboratory protocols.

### Statistical analysis

The effect of diet was tested using one way analysis of variance (ANOVA). The results for initial and final fish weights were subjected to statistical analysis by one-way analysis of variance. Where significant differences at the 5% probability level (p ≤ 0.05) were found between dietary treatments, the means were compared using Duncan’s Multiple Range Test. The data for carcass composition were based on pooled samples from each tank. Pooled Standard Errors of pooled means (± SEM) were calculated to identify the range of means as described by Standard Error **=** s/ √n (where: **s =** sample standard deviation **n =** sample size)

## Results

### Growth performance

The results of this trial show the growth of rainbow trout were affected by the different dietary treatments due to protein substitution and also the supplementation of essential amino acids (Table 6). The control (animal protein blend/fishmeal protein) reference diet fed trout (Ref) gave the superior and significant (P<0.05) performance with a 146% overall increase in body weight in the experimental duration. Conversely, FFS fed trout containing full fat soybean meal was the most inferior amongst the experimental diets with only a 78% increase in weight gain over the same period. Supplementation of this basic formulation with a mixture of essential amino acids in crystalline form resulted in a moderate but significant (P<0.05) improvement in overall growth performance (86.80%) and in particular for MGM+EAAS fed trout when in protected coated form gave an SGR of 125% (P<0.05). The other associated biometric values such as daily live weight gain and protein intake reflected these trends and significance.

**Table 6.** Growth performance and feed utilisation of rainbow trout (Oncorhynchus mykiss) over eight weeks.

| Parameter | Ref diet | FFS | FFS+ EAAS | MGM | MGM+EAAS | MGM+EAAS coated | SEM± |
| --- | --- | --- | --- | --- | --- | --- | --- |
| Initial mean weight (g) | 31.53 | 28.88 | 30.33 | 30.52 | 29.52 | 29.53 | 0.16 |
| Final mean weight (g) | 77.44<sup>a</sup> | 51.48<sup>e</sup> | 56.67<sup>d</sup> | 63.00<sup>c</sup> | 65.95<sup>bc</sup> | 66.48<sup>b</sup> | 1.49 |
| % Increasing body weight | 145.60<sup>a</sup> | 78.30<sup>e</sup> | 86.80<sup>d</sup> | 106.40<sup>c</sup> | 123.40<sup>b</sup> | 125.10<sup>b</sup> | 4.23 |
| SGR %* | 1.61<sup>a</sup> | 1.04<sup>d</sup> | 1.13<sup>d</sup> | 1.29<sup>c</sup> | 1.43<sup>b</sup> | 1.45<sup>b</sup> | 0.04 |
| Feed intake (mg/fish/day) | 861<sup>a</sup> | 680<sup>d</sup> | 740<sup>c</sup> | 770<sup>b</sup> | 780<sup>b</sup> | 760<sup>b</sup> | 9.82 |
| Live wet gain (mg<sup>-1</sup> day<sup>-1</sup>) | 820<sup>a</sup> | 390<sup>e</sup> | 471<sup>d</sup> | 580<sup>c</sup> | 650<sup>b</sup> | 660<sup>b</sup> | 25.33 |
| FCR<sup>a</sup> | 1.05<sup>e</sup> | 1.74<sup>a</sup> | 1.57<sup>b</sup> | 1.33<sup>c</sup> | 1.20<sup>d</sup> | 1.15<sup>d</sup> | 0.04 |
| PER<sup>b</sup> | 2.05<sup>a</sup> | 1.30<sup>f</sup> | 1.47<sup>e</sup> | 1.68<sup>d</sup> | 1.89<sup>c</sup> | 1.94<sup>b</sup> | 0.05 |
| Protein intake (mg<sup>-1</sup> fish<sup>-1</sup> day<sup>-1</sup>) | 402<sup>a</sup> | 300<sup>b</sup> | 321<sup>b</sup> | 345<sup>b</sup> | 344<sup>b</sup> | 341<sup>b</sup> | 22.38 |
| Protein retention (mg<sup>-1</sup> fish<sup>-1</sup> day<sup>-1</sup>) | 140.05<sup>a</sup> | 70.90<sup>d</sup> | 76.00<sup>d</sup> | 91.30<sup>c</sup> | 100.5<sup>c</sup> | 118.10<sup>b</sup> | 5.98 |
| Net protein utilisation (%) | 35.95<sup>c</sup> | 23.64<sup>a</sup> | 23.66<sup>a</sup> | 26.46<sup>a</sup> | 29.21<sup>b</sup> | 34.62<sup>c</sup> | 0.89 |

\*Specific growth rate, <sup>a</sup>Feed Conversion Ratio, <sup>b</sup>Protein Efficiency Ratio) Mean values in rows with different superscripts are significantly different ( $p < 0.05$ )

**Table 7.** Carcass composition (Proximate analysis % whole wet weight)

| Nutrient | Initial<sup>+</sup> | Ref diet | FFS | FFS+ EAAS | MGM | MGM+EAAS | MGM+EAAS coated | SEM±* |
| --- | --- | --- | --- | --- | --- | --- | --- | --- |
| Moisture | 75.08 | 72.27 | 73.07 | 73.61 | 71.99 | 72.24 | 71.94 | 0.67 |
| Protein | 13.29 | 15.87 | 15.17 | 14.62 | 14.55 | 14.48 | 15.85 | 0.64 |
| Lipid | 10.10 | 8.60 | 8.99 | 8.98 | 10.59 | 10.16 | 9.08 | 0.78 |
| Ash | 2.18 | 2.71 | 2.23 | 2.44 | 2.35 | 2.57 | 2.75 | 0.17 |

<sup>+</sup>Composition of initial sample of fish removed at start of experiment. Mean values without superscripts were not significantly different ( $p > 0.05$ ) \*SEM pooled standard error of mean

### Feed utilization

The feed utilisation parameters supported the growth performance metrics above and are presented in Table 6. The feed conversion ratio FCR was better for the Ref diet fed trout (1.05) and poorer for FFS (1.74) (P<0.05). There was an improvement for the FFS+EAAS group. However, all diets based on maize gluten meal displayed very good FCRs with a trend towards improved feed conversion in diets supplemented with both standard and protected amino acid fortification (MGM+EAAS and MGM+EAAS coated. The Net Protein Retention (NPU) in trout receiving each diet gave a similar trend in terms of both growth and feed performance as with the Protein Efficiency Ratio (PER). However, it should be noted that NPU being a direct measurement of protein assimilation efficiency compared to PER which is based directly on protein to total biomass conversion and not protein gain alone. Ref diet fed trout attained 36% ANU and FFS and FFS+EAAS were 23.6% as compared with those fish fed diets MGM, MGM+EAAS and MGM+EAAS coated giving 26.46%, 29.21%, and 34.62% ANUs respectively (P<0.05). In general, plant dietary treatments (especially MGM) supplemented with standard and protected type of amino acids produced significantly (p<0.05) superior dietary protein utilisation.

### True Digestibility Coefficients for essential amino acids

The true digestibility coefficients are displayed in Table 5 for rainbow trout fed the experimental diets with corrections for endogenous protein loss. It is obvious that there was a broad uniformity in the digestibility coefficients reported but a clear enhancement for diets where essential amino acids were added and in their coated forms. These true digestibility coefficients were used to present the digestible amino acid profile for the diets to complement the data for protein bioavailability in real terms compared to the apparent digestibility data more often presented in similar work with fish. It was observed that coated forms of tryptophan, arginine and histidine gave superior and significant (P<0.05) improvements in coefficients in true digestibility over the standard EAA forms. All forms of supplemental essential amino acids in diets for rainbow trout yielded better digestibility values compared to the digestibility of comparable amino acids in the high plant diets and in some cases restored the coefficients to values found for trout fed the reference diet. In general coefficients ranged from values of 82% to most at 88-90%. However, some very low values were recorded such as leucine (52%) and tryptophan and threonine in the full fat soya treatment (70 and 71%) respectively indicating poor digestibility for this ingredient in un-supplemented EAAs.

### Carcass composition

On the basis of carcass composition (Table 7) the composition of the rainbow trout did not vary significantly (P>0.05) when assessed at the termination of the study after 8-weeks on the experimental diets. However, it is interesting that the Ref diet control group and MGM+EAAS coated diets produced the highest protein carcass levels in rainbow trout. This agreed with the better protein utilisation NPUs values observed for these fish.

## Discussion

This investigation on optimising diets for rainbow trout using higher plant based protein ingredients was justified to meet the global need for sustainability facing the aquaculture industry. This is now a priority research area to test various oilseed and pulses and cereal derived concentrates as overviewed by Gatlin *et al*. (2007). The reduction in fishmeal inclusion in contemporary diets places pressure to obtain sufficient balance to ensure the correct essential amino acid profile in commercial feed formulations. Our experiment showed the viability of successfully using coated crystalline amino acids to effectively restore the limitations of utilising maize gluten protein where lysine and arginine were included at their higher substitution level of fishmeal compared to the reference formulation diet. For example, a marked improvement was observed for maize gluten meal because lysine is a well-known first limiting essential amino acid. Its deficiency can compromise the ideal essential amino acid profile in livestock and is highly significant in swine nutrition when diets are high in this ingredient relative to soybean meal (Boisen et al. (2000). This is similar for many fish and other animal studies where maize products have been utilised in formulated commercial diets and under experimental conditions with reference to NRC (2011).

Previous work has identified the influence of multiple amino acid supplementation on improving performance and feed utilisation in rainbow trout (Davies and Morris, 1997). In this study, a series of soyabean containing diets replacing up to 66% of fishmeal were then complemented with crystalline amino acids. namely, methionine only, dual supplemented with two methionine and lysine levels and finally, a group complex comprising methionine, lysine, tryptophan, threonine, arginine and histidine. The results indicated that soyabean meal (SBM) was inferior to the reference fishmeal protein when SBM was used alone. No restoration in growth, feed efficiency and apparent net protein utilisation was obtained by either methionine only or with both methionine and lysine supplementation. However, multiple amino acid addition to the diet yielded an improved percentage weight gain, specific growth rate and a small improvement in apparent net protein utilisation for rainbow trout. Other work on rainbow trout was closer in concept to the current study (Davies, Morris and Baker, 1997) investigating the addition of crystalline lysine to correct its deficiency in wheat gluten meal protein. Wheat gluten meal has been used in salmonid diets but is constrained by lysine being a first limiting essential amino acid. This study with maize gluten meal extends this work further for a more generic product in rainbow trout diet formulations and reinforces its use with protected essential amino acid fortification.

The supplementation of diets for pigs and poultry has long been deemed important in flexible diet formulations where corn or maize gluten meals have been the basis of diets with lower levels of soybean and animal byproducts. Precision diets using more accurate nutrient data and for digestibility of individual amino acids and protein form the basis of such feeds using ileal digestibility data and coupled with growth trials in younger animals (piglets and chicks) as well as later production stages. It’s the fast growing juvenile stages, where protein accretion is much more critically dependent on essential amino acid supply and balance to meet the ‘ideal protein’ underpins the concept first proposed by Wang and Fuller (Wang and Fuller, 1989; D’Mello, 1993) in swine. The most important single factor affecting the efficiency of protein utilisation is the profile of digestible essential amino acids entering the small intestine. Assuming a constant ideal amino acid profile of absorbed protein where the requirements of all amino acids can be calculated when the relative requirement of one individual amino acid (usually lysine) has been determined. This principle holds true in the current investigation with rainbow trout. However, a faecal stripping method was applied to obtain faecal material, and this cannot be directly compared to total faecal collection protocols used in many animal species. Previous researchers (Boisen, Hvelplund and Weisbjerg, 2000) have used the amino acid deletion method to arrive at an estimate of optimum dietary EAA pattern for rainbow trout. With 40% deletions of a single EAA from the control pattern. Based on N utilisation data, an estimate of optimum dietary EAA pattern for rainbow trout can be constructed. Using a linear formulation software, we compared the dietary EAA pattern for rainbow trout, based on 1) amino acid composition of rainbow trout whole-body protein, 2) EAA requirements for rainbow trout published by the National Research Council (NRC, 2011).

Modern approaches to undertake precision diet formulations for fish must depend on accurate digestibility data being obtained and this has been fraught with technical difficulties compared to terrestrial animal studies and is stated in previous work for rainbow trout to evaluate minimize nitrogen excretion and maximize nitrogen retention as in our current study (Green and Hardy, 2002). However, the present investigation featured a separate trial to determine the intestinal endogenous amino acid excretion with the use of a low-protein diet. This diet was carefully formulated to contain 2% crude protein from casein as a purified diet. Prior (unpublished) data by the author that a low rather than a zero protein diet prevents over stimulation of endogenous protein due to gut enterocyte sloughing into the digesta resulting in over estimation of nitrogen losses. Assuming that residual amino acids supplied in a low-protein diet should be subtracted from the exogenous dietary derived faecal source, the true amino acid excretion was calculated. To our knowledge this current study to assess essential amino acid supplementation with fishmeal being replaced by soybean meal and maize gluten is unusual in providing true amino acid digestibility data for trout in combination with supplementary essential amino acids. This study adds true digestibility data for plant-based trout diets, complementing earlier methods.

A comprehensive review of fishmeal replacement by various alternative ingredients on growth performance of fish and adoptability has been recently published (Macusi *et al*., 2023) and for aquaculture sustainability as reported for Nile tilapia to examine true and apparent digestibility of protein and amino acids in feed (Ribeiro *et al*., 2011) Hussain *et al*., 2024).

As such, the practical applications of crystalline essential amino acids for both shrimp and fish diets are now gaining potential within the industry for maximum efficacy of plant ingredients and for enhanced performance, welfare and cost of production shown by Nunes *et al*. (2014) and Adhikari et al. (2025). With the emerging use of closed Recirculation Aquaculture Systems (RAS) for fish rearing there is an increased interest for low fishmeal formulated diets to meet the intensive production of salmonid species as recently stated by Fanizza *et al*. (2023). This will require the establishment of optimal use of protein and supplemental essential amino acids to improve nutrient digestibility and environmental impact of nitrogen excretion. The present study provides data for rainbow trout that can be useful to meet such concepts in practice.

Some investigations have been done to make economic savings by attempting to reduce the crude protein levels of diets by supplementation of the first, secondary and tertiary essential amino acids in advanced feeds for salmon, trout and for other fish species such as tilapia (Bell and Davies, 2024). Also, for channel catfish, *Ictalurus punctatus* investigated successfully by Salem *et al*. (2022). In this way the ‘ideal’ protein ratio and quantitative levels of essential amino acids can be restored to some effect. The optimum protein to energy ratio of salmonid diets has been well established but there is more scope for refinement by lowering the protein levels in diets. This can lead to higher N and amino acid retention with less metabolic, physiologic and less environmental impact into the aquatic medium (Lee *et al*., 2019).

The current investigation using rainbow trout as our salmonid model has given evidence that crystalline amino acids may impart a capacity to improve protein utilisation of diets formulated with higher plant ingredient levels and offset to some degree the essential amino acids discrepancy encountered. The ideal amino acid requirements for meeting the needs of alternative protein sources for complete diet formulations needs to be validated for different phases of production and for many more potential candidate ingredients such as various pulses and legumes and cereal derived proteins beyond soybean and maize gluten meals. With ingredients like SCPs and insect meals we need more precise digestibility data for different fish species if we are to correct any essential amino acid deficiencies using various crystalline forms in the diet.

## Conclusion

It was demonstrated that partial replacement of fishmeal with soybean meal and maize gluten meals could be achieved without serious impairment of growth performance and feed utilisation in trout, but higher level of inclusion necessitated crystalline amino supplementation to mitigate nutritional constraints. Although the reference diet fed trout produced the better overall growth performance and protein retention, plant ingredients like soybean and maize could become more effective in linear least cost formulations for salmonid fish species like trout and salmon provided essential amino acids are provisioned. Meeting the circular economy with plant byproducts will necessitate more refined data on digestibility and potential for EAAs for precision diet formulations. The cost benefits of using synthetic and crystalline essential amino acids must also be factorized in modern feed formulations to meet the sustainability agenda towards advancing the aquaculture sector.

